## Appendix for "Non-canonical contribution of human Dicer helicase domain in antiviral innate immune response"

**Table of Contents**


| | Page |
|---|---|
| **Appendix Figure S1** | 2 |
| **Appendix Figure S2** | 3 |
| **Appendix Figure S3** | 4 |
| **Appendix Figure S4** | 5 |
| **Appendix Table S1** | 6 |

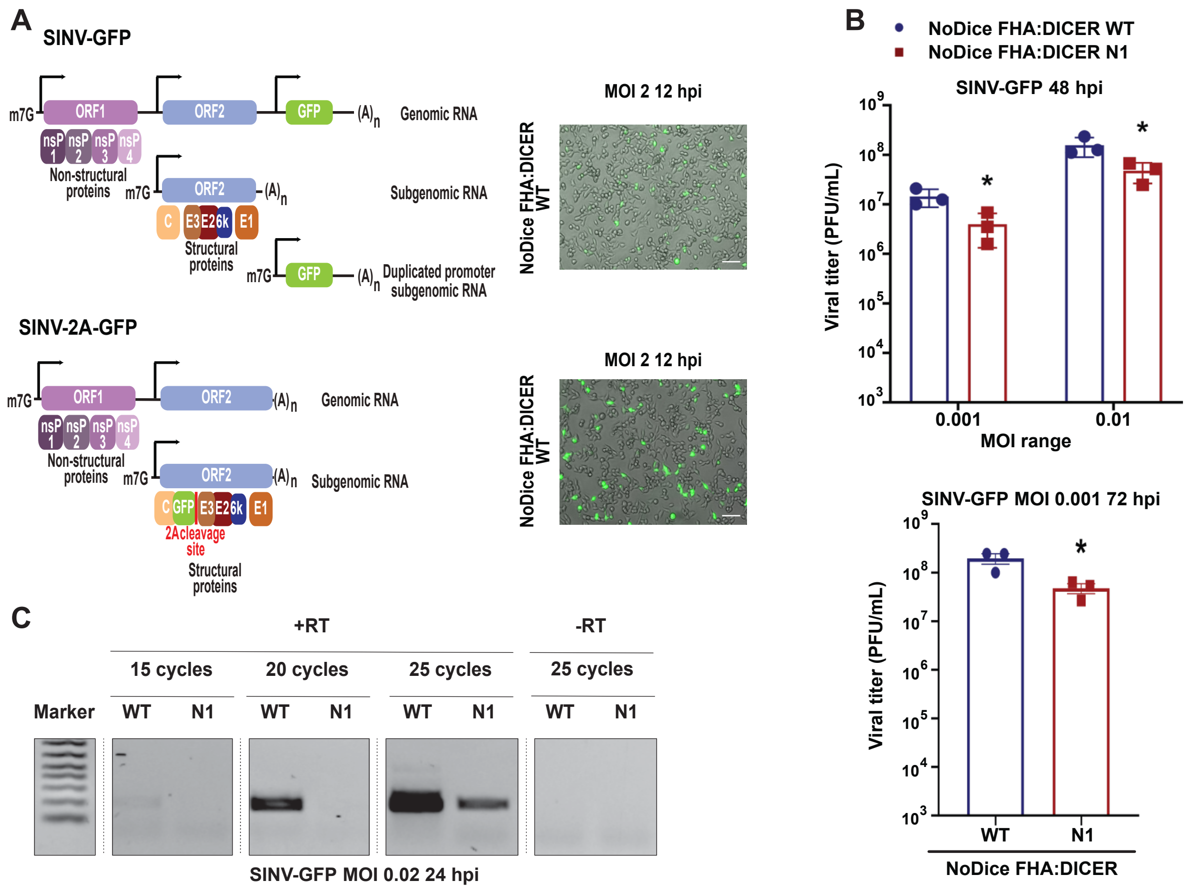


**Appendix Figure S1.** Dicer N1 reduces SINV replication at longer infection times and decreases antigenome production.

**A.** Schematic representation of SINV-GFP and SINV-2A-GFP genome organization and representative fluorescent pictures of GFP expressed by the two viruses upon infection of NoDice FHA:DICER WT cells at an MOI of 2 for 12h. Scale bar = 50µm, (n = 3 biological replicates). ORF: Open reading frame, nsP: non-structural proteins, C: capsid, E1 to E3: envelope proteins.

**B.** Mean (+/- SEM) SINV-GFP viral titer in NoDice FHA:DICER WT and N1 infected at an MOI 0.001 and 0.01 for 48h (up) and 0.001 for 72h (bottom) from plaque assay quantification (n = 3 biological replicates). Upper panel: ordinary two-way ANOVA test with Sidak's correction. *: $p < 0.05$. Bottom panel: Unpaired t-test. *: $p<0.05$.

**C.** Agarose gel of RT-PCR on SINV antigenome in SINV-GFP infected NoDice FHA:DICER WT and N1 cells at an MOI of 0.02 for 24h (n = 3 biological replicates).

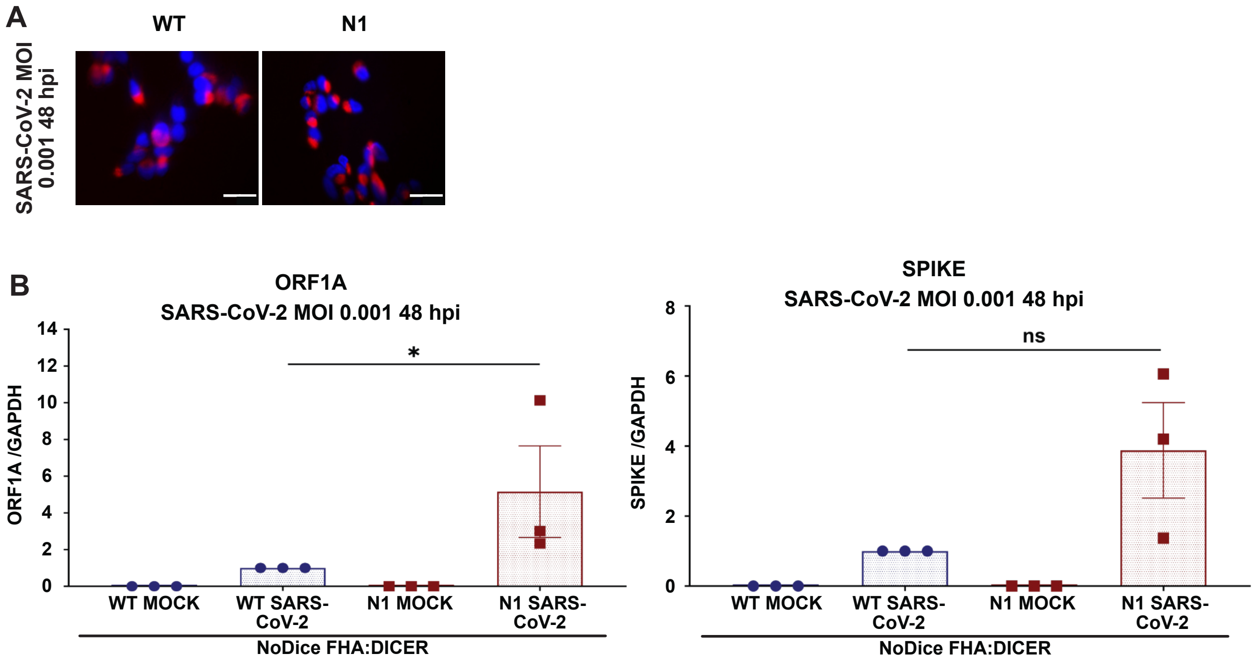


**Appendix Figure S2.** Dicer N1 enhances SARS-CoV-2 replication.
**A.** Immunofluorescence analysis on NoDice FHA:DICER WT and N1 cells in SARS-CoV-2 infected cells at an MOI of 0.001 for 48h. J2 antibody (red) was used to detect dsRNA upon infection. DAPI was used to stain the nuclei (blue). Scale bar = 100 µm, (n = 3 biological replicates).
**B.** RT-qPCR on SARS-CoV-2 genome (ORF1A and SPIKE) in NoDice FHA:DICER WT and N1 cells infected with SARS-CoV-2 at an MOI of 0.001 for 48h. Mean (+/-SEM); n = 3 biological replicates. Unpaired t-test. *: $p<0.05$; ns: non-significant.

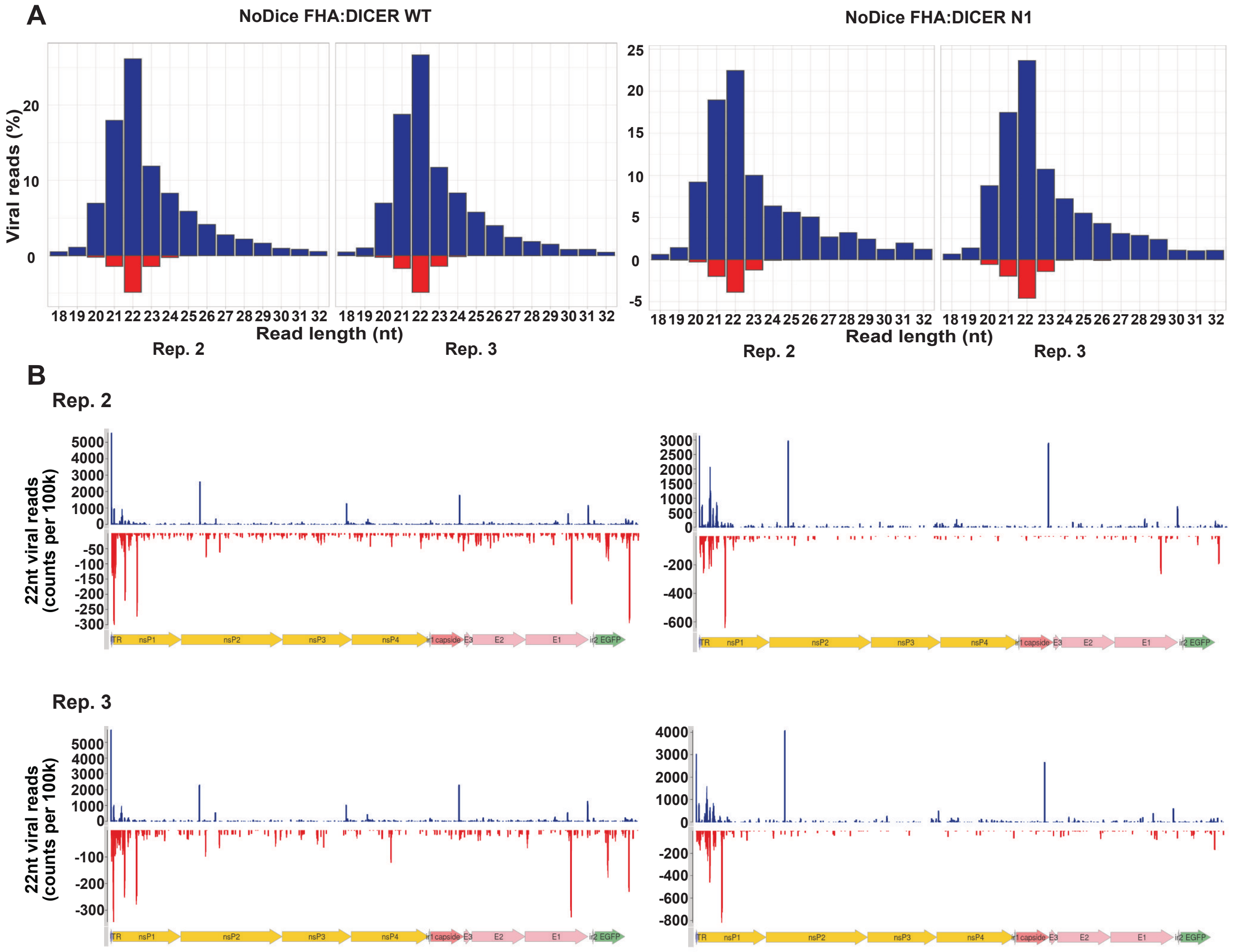


**Appendix Figure S3.** Dicer N1 cells do not accumulate more virus-derived small RNAs than Dicer WT cells.

**A.** Representative histograms of the distribution of viral reads (in percent) per RNA length upon small RNA sequencing for replicates 2 and 3 in NoDice FHA:DICER WT (left) and N1 (right) cells infected with SINV-GFP at an MOI of 0.02 for 24h. Blue: positive strand; red: negative strand.

**B.** Representative graphs of the mapping of 22-nt-reads on SINV-GFP genome for replicates 2 and 3 in NoDice FHA:DICER WT (left) and N1 (right) cells.

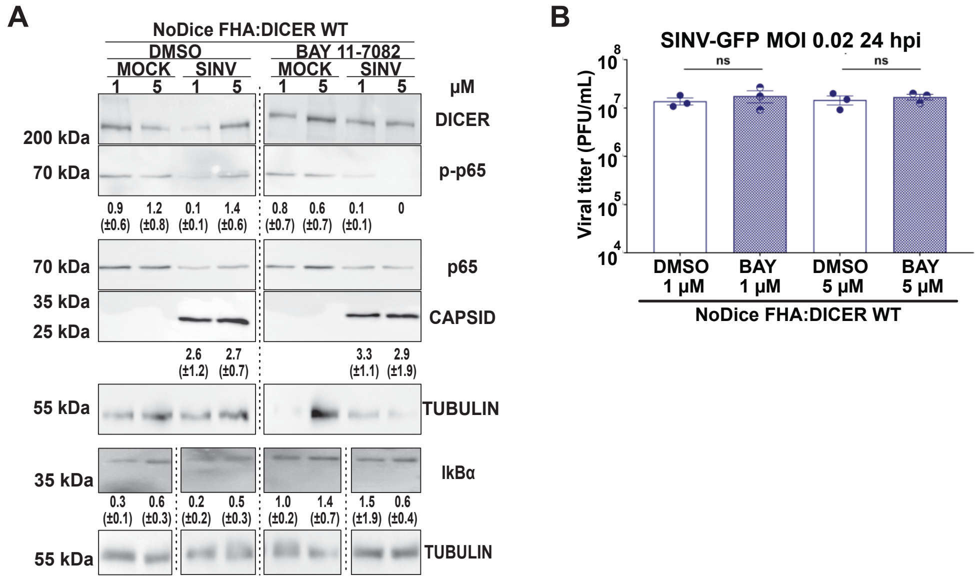


**Appendix Figure S4.** NF-kB/p65 is not involved in the infection outcome in Dicer WT cells.
**A.** Western blot analysis of DICER, p-p65, p65, CAPSID and IkBα expression in SINV-GFP infected NoDice FHA:DICER WT cells at an MOI of 0.02 for 24h. Before infection, cells were treated with the NF-kB/p65 inhibitor, BAY 11-7082 or the vehicle (DMSO) at the indicated concentrations for 1 h. Alpha-Tubulin was used as loading control. Mean (+/- SD) of bands quantification of three independent replicates normalized to Tubulin are represented under the corresponding lane; p-p65 = p-p65/p65 quantification.
**B.** Mean (+/- SEM) of SINV-GFP viral titers in the same samples as in 1, infected at an MOI of 0.02 for 24h (n = 3 biological replicates) from plaque assay quantification. Ordinary one-way ANOVA test with Sidak's correction. ns: non-significant.

**Appendix Table S1.** List of oligonucleotides used in this study.

| Type | Target | Fw | Rev |
|---|---|---|---|
| **PCR primers for mutagenesis** | PKR K296R | CGTTATTAGACGTGTTAAATATAATAACGAGAAGG | ACACGTCTAATAACGTAAGTCTTTCCG |
| | PKR T451A | TAAGGGAGCATTGCGATACATGAGCCCAGAAC | CGCAATGCTCCCTTACTCCTTGTTCGCTTTCC |
| | MYC EMPTY | AGGATCTGACCCAGCTTTCTTGTACAAAGTGG | GCTGGGTCAGATCCTCTTCTGAGATGAGTTT |
| | N1-CM | ACCAATTCAGTCGACATGGAAGCAGAATTC | GAATTCTGCTTCCATGTCGACTGAATTGGT |
| | ΔHEL1 | CGCCAAGAAAATATCAGGTTAGTAATGCTGAAACTGC | GTTGCAGTTTCAGCATTACTAACCTGATATTTTCTTG |
| | ΔHEL2i | TTAGACAGAAATTTTCCTTCTCCTTTTACCAAC | AAAATTTCTGTCTAAGACCACCAGGTCAG |
| | ΔHEL2 | CAGAGACACCTGTCATGGATGATGATCACG | TGACAGGTGTCTCTGGCTTCTCTTTTTCTTC |
| **RT-PCR primers** | RT specific SINV | ATTGACGGCGTAGTACACACTATTGAATCAAACAGCC GACCA | / |
| | PCR SINV antigenome | TAGACGTAGACCCCCAGAGTC | CATTCTACGAGCCGGTGCGC |
| **qPCR primers** | SINV genome | CCACTACGCAAGCAGAGACG | AGTGCCCAGGGCCTGTGTCCG |
| | GAPDH | CTTTGGTATCGTGGAAGGACT | CCAGTGAGCTTCCCGTTCAG |
| | OAS3 | TGCTGCCAGCCTTTGACGCC | TCGCCCGCATTGCTGTAGCTG |
| | PTGS2 | ACCCACTCCAAACACAGTGC | GCTTCCCAGCTTTTGTAGCC |
| | APOBEC3B | GACCCTTTGGTCCTTCGAC | GCACAGCCCCAGGAGAAG |
| | MX1 | AAGCTGATCCGCCTCCACTT | TGCAATGCACCCCTGTATACC |
| | IFN ß | AAGGCCAAGGAGTACAGTC | ATCTTCAGTTTVGGAGGTAA |
| | IFIT3 | ATGAGTGAGGTCACCAAGAATTCCC | GGCTGCCTCGTTGTTACCAT |
| **Northern blot probes** | miR-16 | CGCCAATATTTACGTGCTGCTA | |
| | U6 | GCAGGGGCCATGCTAATCTTCTCTGTATCG | |
